# Decoding the Neural Signatures of Valence and Arousal From Portable EEG Headset

**DOI:** 10.1101/2021.07.23.453533

**Authors:** Nikhil Garg, Rohit Garg, Apoorv Anand, Veeky Baths

## Abstract

Emotion classification using electroencephalography (EEG) data and machine learning techniques has been on the rise in the recent past. However, past studies uses data from medical-grade EEG setup with long set-up time and environment constraints. This paper focuses on classifying emotions on the valence-arousal plane using various feature extraction, feature selection and machine learning techniques. We evaluate different feature extraction and selection techniques and propose the optimal set of features and electrodes for emotion recognition. The images from the OASIS image dataset were used to elicit valence and arousal emotions, and the EEG data was recorded using the Emotiv Epoc X mobile EEG headset. The analysis is carried out on publicly available datasets: DEAP and DREAMER for benchmarking. We propose a novel feature ranking technique and incremental learning approach to analyze performance dependence on the number of participants. Leave-one-subject-out cross-validation was carried out to identify subject bias in emotion elicitation patterns. The importance of different electrode locations was calculated, which could be used for designing a headset for emotion recognition. The collected dataset and pipeline are also published. Our study achieved a root mean square score (RMSE) of 0.905 on DREAMER, 1.902 on DEAP, and 2.728 on our dataset for valence label and a score of 0.749 on DREAMER, 1.769 on DEAP and 2.3 on our proposed dataset for arousal label respectively.

## 1 INTRODUCTION

The role of human emotion in cognition is vital and has been studied for a long time, with different experimental and behavioural paradigms. Psychology researchers have tried to understand human perception through surveys for a long time. Recently, with the increasing need to learn about human perception, without human biases and conception of various emotions across people (Ekman, 1972), we observe the increasing popularity of neurophysiological recordings and brain imaging methods. Since emotions are triggered almost instantly, Electroencephalography (EEG) is an attractive choice due to its better temporal resolutions and mobile recording devices. (Tuncer et al., 2021; Lang, 1995; Katsigiannis and Ramzan, 2018; Koelstra et al., 2012; Ko et al., 2021; Moss et al., 2003).

However, most pattern recognition benchmarks for decoding human emotions from EEG signals have been performed with research-grade EEG recording systems with large setup times, sophisticated recording setup, and cost. Although a portable EEG headset has a lesser signal-to-noise ratio, its low-cost and easy use makes it an attractive choice for collecting data from a wider population sample and overcoming the problem of insufficient uniform EEG data for algorithmic research. The algorithmic pipeline of decoding user intentions through neurophysiological signals consists of denoising, pre-processing, feature extraction, electrode and feature selection, and classification. Although there are deep-learning algorithms (Haselsteiner and Pfurtscheller, 2000; Jeevan et al., 2019; Karlekar et al., 2018; Mahajan and Baths, 2021; Schirrmeister et al., 2017; Übeyli, 2009; Zhou et al., 2018; Jin and Kim, 2020; Tao et al., 2020) which claim to do the frequency decomposition, feature extraction, and classifier training in the hidden layers, their explainability is limited, and amount of training data required is huge. Machine learning-based emotion recognition hence performs weighted Spatio-temporal averaging of EEG signals. While several feature extraction methods were reported in the past, it is crucial to understand which methods are suited for emotion recognition and optimize the set of features for performance. Moreover, the electrodes’ relative importance can help explain the significance of different regions for emotion elicitation. This could, in turn, help in optimizing the electrode locations while conducting EEG-based studies.

In this study, first, we propose a protocol for eliciting emotions by presenting selected images from the OASIS dataset (Kurdi et al., 2016) and signal recording through a low-cost, portable EEG headset. Second, we create a pipeline of pre-preprocessing, feature extraction, electrode and feature selection, classifier for emotional response (Valence and Arousal) decoding and evaluate it for our dataset and two open-source datasets; incremental training to demonstrate the dependence of performance on population sample size is presented. Third, we rank different categories of feature extraction techniques to evaluate the applicability of feature extraction techniques for highlighting the patterns indicative of emotional response. Moreover, we analyze the electrode importance and rank different brain regions for their importance. Fourth, we ask if we can automate the feature selection and electrode selection techniques for BCI pipeline engineering and validate the procedure with a qualitative and quantitative comparison with neuroscience literature. Lastly, we publish the proposed pipeline and recorded dataset for community.

In the past, the scope of using electrophysiological data for emotion prediction has widened and led to standardized 2D emotion metrics of valence and arousal (Russell, 1980) to train and evaluate pattern recognition algorithms. Human brain-recording experiments have been conducted as part of effort to associate emotion quantitatively with words, pictures, sounds, and videos (Moors et al., 2013; Mohammad, 2018; Leite et al., 2012; Lane et al., 1999; Gerber et al., 2008; Warriner et al., 2013; Eerola and Vuoskoski, 2011; Lang, 1995; Kurdi et al., 2016). EEG frequency band has been found to be dominant during different roles, corresponding to various emotional and cognitive states (Klimesch, 2012, 1999; Klimesch et al., 1990; Klimesch, 1996; Kamiński et al., 2012; Bauer et al., 2007; Berens et al., 2008; Jia and Kohn, 2011). Besides using energy spectral values, researchers use many other features such as frontal asymmetry, differential entropy and indexes for attention, approach motivation and memory. “Approach” emotions such as happiness are associated with left hemisphere brain activity, whereas “withdrawal,” such as disgust, emotions are associated with right hemisphere brain activity (Coan et al., 2001; Davidson et al., 1990). The left to right alpha activity is therefore used for approach motivation. The occipito-parietal alpha power has been found to have correlations with attention(Misselhorn et al., 2019; Smith and Gevins, 2004). Fronto-central increase in theta and gamma activities has been proven essential for memory-related cognitive functions(Shestyuk et al., 2019). Differential entropy combined with asymmetry gives out features such as differential and rational asymmetry for EEG segments are some recent developments as forward-fed features for neural networks(Duan et al., 2013; Torres P. et al., 2020).

In an attempt to classify emotions using EEG signals, many time-domain, frequency-domain, continuity, complexity(Gao et al., 2019; Galvão et al., 2021), statistical, microstate (Milz et al., 2016; Lehmann, 1990; Shen et al., 2020b), wavelet-based (Jie et al., 2014). Empirical features (Subasi et al., 2021; Patil et al., 2019) have been used to aid better classification results using advanced ensemble learning techniques (Fang et al., 2021) or using deep networks, often referred to as bag of deep features (Asghar et al., 2019). We have summarized the latest studies using EEG to recognize the emotional state in Table **??**

This paper is organized as follows. Section 2 provides description of the three datasets used for our analysis. The theoretical background and the details of pre-processing steps (referencing, filtering, motion artifact and rejection and repair of bad trials) are discussed in section 3. Section 4 addresses the feature extraction details and provides overview of features extracted. Section 5 describes the feature selection procedure adapted in this work. Section 6 presents our experiments and results. This is followed by section 7 for discussion of experiments performed and results obtained in this work. Finally, Section 8 summarizes the conclusion and future scope of this work.

## MATERIALS AND METHODS

### 2 DATASETS

#### 2.1 OASIS EEG dataset

##### 2.1.1 Stimuli selection

The OASIS image dataset (Kurdi et al., 2016) consists of a total 900 images from various categories such as natural locations, people, events, inanimate objects with various valence and arousal elicitation values. Out of 900 images, 40 images were selected to cover the whole spectrum of valence and arousal ratings as shown in Fig 1.

**Figure 1.**
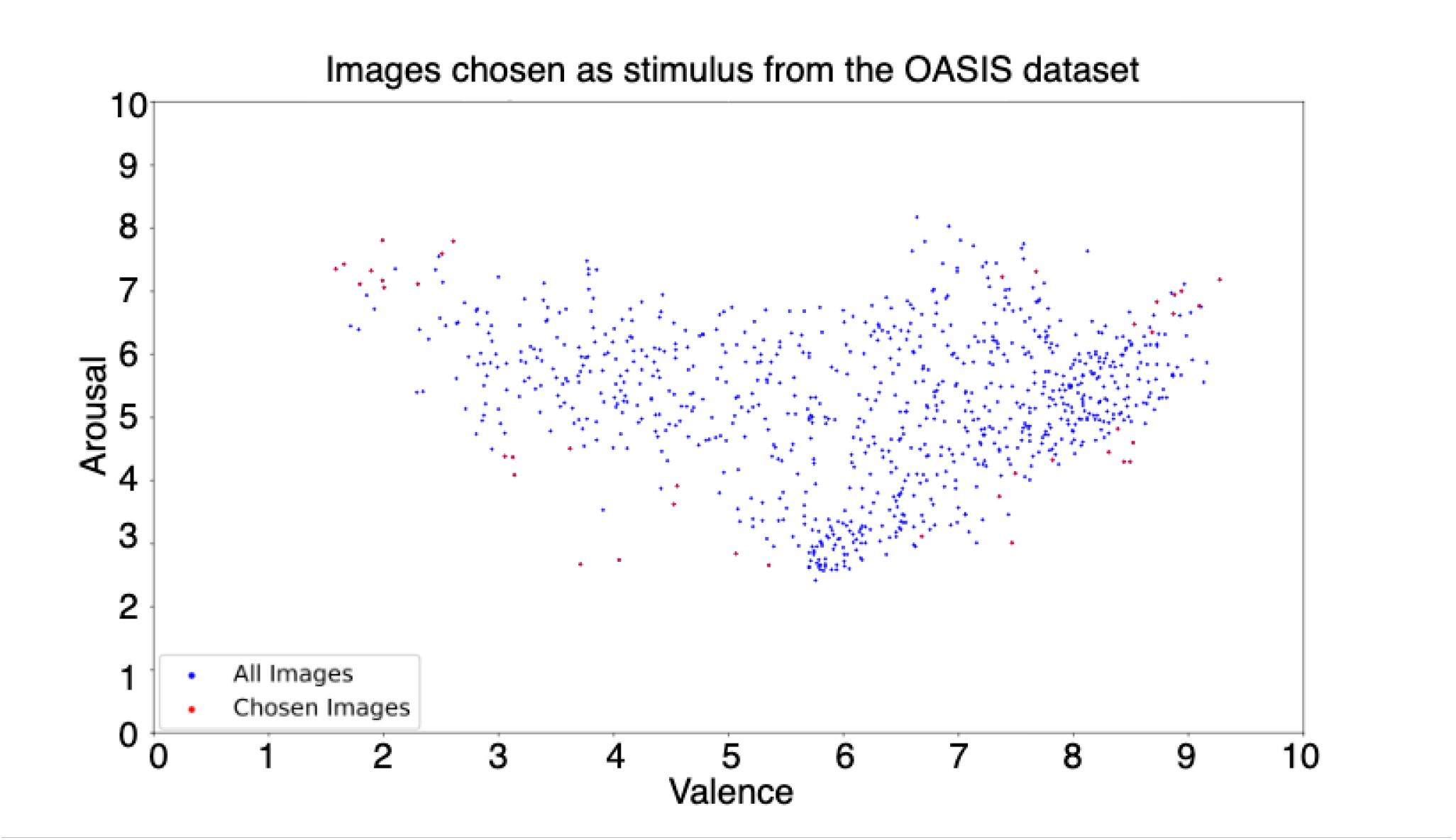
Valence and arousal ratings of OASIS dataset. Valence and arousal ratings of entire OASIS (Kurdi et al., 2016) image dataset (blue) and of the images selected for our experiment (red). The images were selected to represent each quadrant of the 2D space.

##### 2.1.2 Participants and device

The experiment was conducted in a closed room with the only source of light being the digital 21” Samsung 1080p monitor. Data was collected from fifteen participants of mean age 22 with ten males and five females using EMOTIV Epoc EEG headset consisting of 14 electrodes according to the 10-20 montage system at a sampling rate of 128Hz and only the EEG data corresponding to the image viewing time were segmented using markers and used for analysis.

The study was approved by the Institutional Ethics Committee of BITS, Pilani (IHEC-40/16-1). All EEG experiments/methods were performed in accordance with the relevant guidelines and regulations as per the Institutional Ethics Committee of BITS, Pilani. All participants were explained the experiment protocol and written consent for recording the EEG data for research purpose was obtained from each of the subject.

##### 2.1.3 Protocol

The subjects were explained the meaning of valence and arousal before the start of the experiment and were seated at a distance of 80-100 cms from the monitor.

The images were shown for 5 seconds through Psychopy (Peirce et al., 2019), and the participants were asked to rate valence and arousal on a scale of 1 to 10 before proceeding to the next image, as shown in Fig 2. The ratings given in the OASIS image dataset were plotted against the ratings reported by the participants in order to draw correlation between them, as shown in Fig 3.

**Figure 2.**
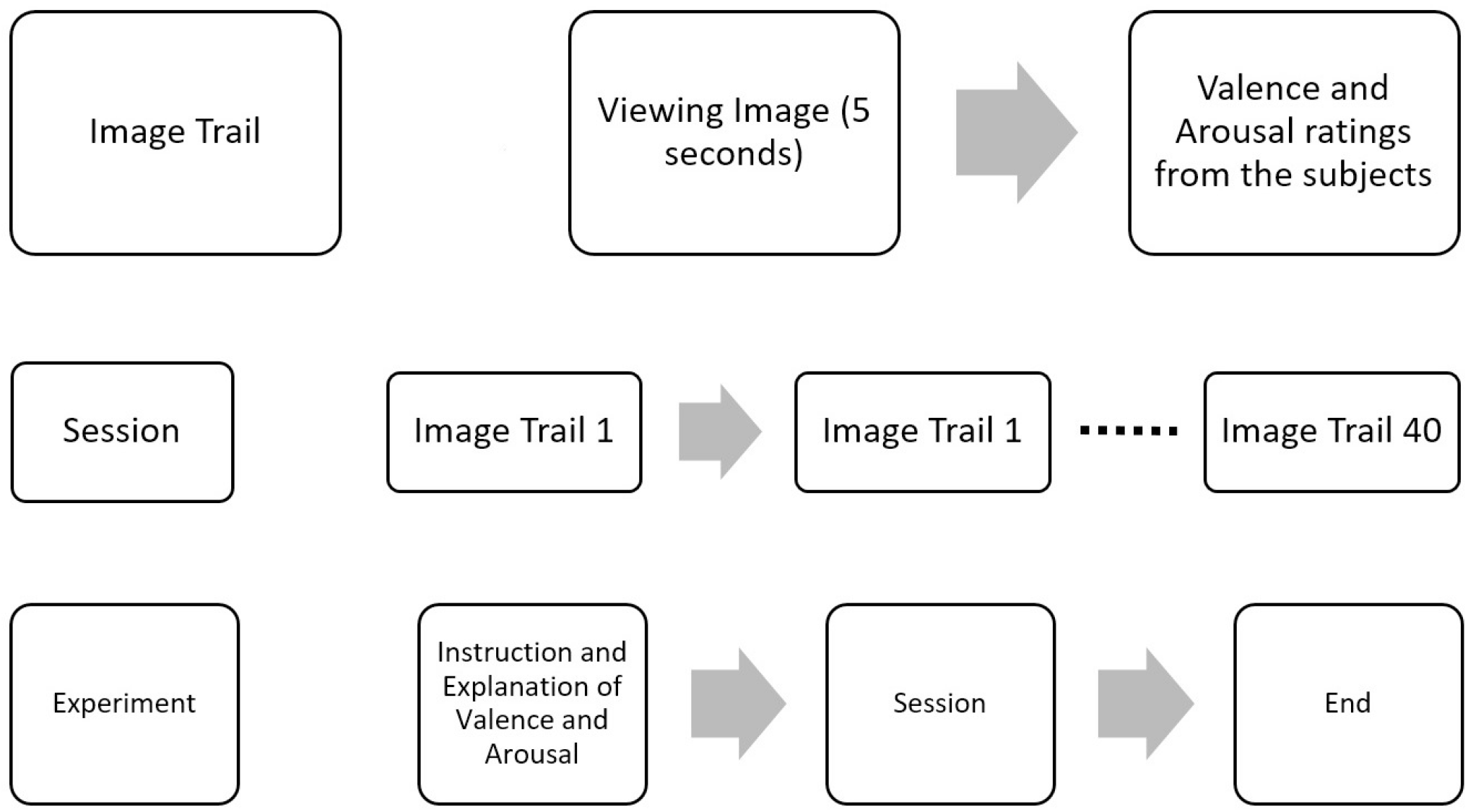
EEG Data collection protocol. Experiment protocol for the collection of EEG data. 40 images from OASIS dataset were shown to elicit emotion in valence and arousal plane. After presenting each image, ratings were collected from participant.

**Figure 3.**
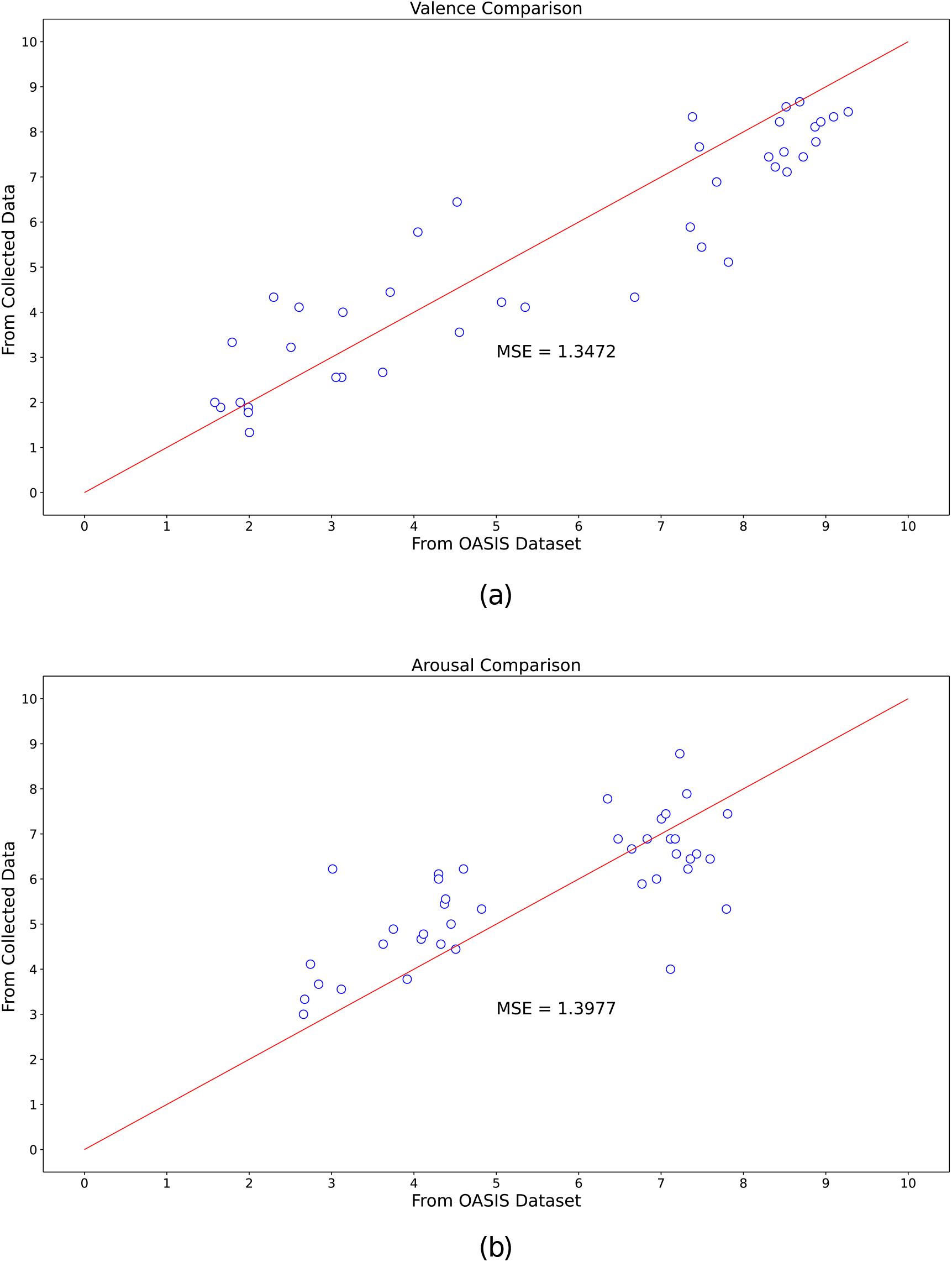
Comparison of actual and self-reported valence and arousal ratings. Valence and arousal ratings reported by the participants during the EEG data collection and ratings from the OASIS image dataset.

#### 2.2 DEAP

DEAP dataset(Koelstra et al., 2012) has 32 subjects; each subject was shown 40 music videos one min long. Participants rated each video in terms of arousal levels, valence, like/dislike, dominance, and familiarity. Data was recorded using 40 EEG electrodes placed according to standard 10-20 montage system. The sampling frequency was 128Hz. In this analysis, we consider only 14 channels (AF3, F7, F3, FC5, T7, P7, O1, O2, P8, T8, FC6, F4, F8, AF4) for the sake of uniformity with other two datasets.

#### 2.3 DREAMER

DREAMER(Katsigiannis and Ramzan, 2018) dataset has 23 subjects; each subject was shown 18 videos at a sampling frequency 128Hz. Audio and visual stimuli in the form of film clips were employed to elicit emotional reactions to the participants of this study and record EEG and ECG data. After viewing each film clip, participants were asked to evaluate their emotion by reporting the felt arousal (ranging from uninterested/bored to excited/alert), valence (ranging from unpleasant/stressed to happy/elated), and dominance. Data was recorded using 14 EEG electrodes.

## 3 PREPROCESSING

Raw EEG signals extracted out of the recording device are continuous, unprocessed signals containing various kinds of noise, artifacts and irrelevant neural activity. Hence, lack of EEG pre-processing can reduce the signal-to-noise ratio and introduce unwanted artifacts into the data. In pre-processing step, noise and artifacts presented in the raw EEG signals are identified and removed to make them suitable for analysis on the further stages of experiment. The following subsections discuss each step of pre-processing (referencing, filtering, motion artifact and rejection and repair of bad trials) in more detail.

### 3.1 Referencing

The average amplitude of all electrodes for a particular time point was calculated and subtracted from the data of all electrodes. This was done for all time points across all trials.

### 3.2 Filtering

A Butterworth bandpass filter of 4^*th*^ order was applied to filter out frequencies between 0.1Hz and 40Hz

### 3.3 Motion Artifact

Motion artifacts can be removed by using Pearson Coefficients (Onikura et al., 2015). The gyroscopic data (accelerometer readings) and EEG data were taken corresponding to each trial. Each of these trials of EEG data was separated into its independent sources using Independent Component Analysis (ICA) algorithm. For the independent sources obtained corresponding to a single trial, Pearson coefficients were calculated between each source signal and each axis of accelerometer data for the corresponding trial. The mean and standard deviations of Pearson coefficients were then calculated for each axis obtained from overall sources. The sources that had Pearson coefficient 2 standard deviations above mean for any one axis were high pass filtered for 3Hz using a Butterworth filter as motion artifacts exist at these frequencies. The corrected sources were then projected back into the original dimensions of the EEG data using the mixing matrix given by ICA.

### 3.4 Rejection and repair of bad trials

Auto Reject is an algorithm developed by Mainak et al. (Jas et al., 2017) for the rejection of bad trials in Magneto-/Electro-encephalography (M/EEG data), using a cross-validation framework to find the optimum peak to peak threshold to reject data.

- We first consider a set of candidate thresholds *ϕ*.
- Given a matrix of dimensions (epochs x channels x time points) by X ∈ R N×P, where N is the number of trials/epochs P is the number of features. P = Q*T, Q being the number of sensors, and T the number of time points per sensor.
- The matrix is split into K folds. Each of the K parts will be considered the training set once, and the rest of the K-1 parts become the test set.
- For each candidate threshold, i.e. for each

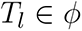

we apply this candidate peak to peak threshold(ptp) to reject trials in training set known as bad trials, and the rest of the trials become the good trials in the training set.

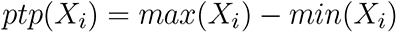

where *X*_*i*_ indicates a particular trial.

- A is the peak to peak threshold of each trial, *G*_*l*_ is the set of trials whose ptp is less than the candidate threshold being considered

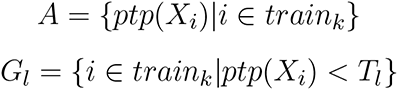
- Then the mean amplitude of the good trials (for each sensor and their corresponding set of time points) is calculated

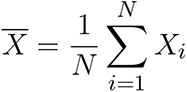
- While the median amplitude of all trials is calculated for the test set 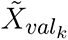
- Now the Frobenius norm is calculated for all K folds giving K errors *e*_*k*_ *∈ E*; mean of all these errors is mapped to the corresponding candidate threshold.

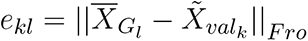
- The following analysis was done taking all channels into consideration at once, thus it is known as auto-reject global
- Similar process can be considered where analysis can be done for each channel independently i.e data matrix becomes(epochs x 1 x time points) known as the local auto-reject, where we get optimum thresholds for each sensor independently.
- The most optimum threshold is the one that gives the least error

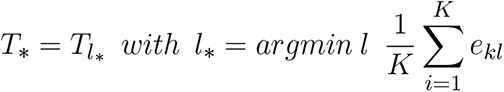

As bad trials were already rejected in the DEAP and DREAMER dataset, we do not perform automatic trial rejection in them.

## 4 FEATURE EXTRACTION

In this work, the following set of 36 features were extracted from the EEG signal data with the help of EEGExtract library (Saba-Sadiya et al., 2020) for all three datasets:

- Shannon Entropy (S.E.)
- Subband Information Quantity for Alpha [8 Hz - 12 Hz], Beta [12 Hz - 30 Hz], Delta [0.5 Hz - 4 Hz], Gamma [30 Hz - 45 Hz] and Theta[4 Hz - 8 Hz] band (S.E.A., S.E.B., S.E.D., S.E.G., S.E.T.)
- Hjorth Mobility (H.M.)
- Hjorth Complexity (H.C.)
- False Nearest Neighbour (F.N.N)
- Differential Asymmetry (D.A., D.B., D.D., D.G., D.T.)
- Rational Asymmetry (R.A., R.B., R.D., R.G., R.T.)
- Median Frequency (M.F.)
- Band Power (B.P.A., B.P.B., B.P.D., B.P.G., B.P.T.)
- Standard Deviation (S.D.)
- Diffuse Slowing (D.S.)
- Spikes (S.K.)
- Sharp spike (S.S.N.)
- Delta Burst after Spike (D.B.A.S.)
- Number of Bursts (N.B.)
- Burst length mean and standard deviation (B.L.M., B.L.S.)
- Number of Suppressions (N.S.)
- Suppression length mean and standard deviation (S.L.M., S.L.S.)

These features were extracted with a 1s sliding window and no overlap. The extracted features can be categorized into two different groups based on the ability to measure the complexity and continuity of the EEG signal. The reader is encouraged to refer to the work done by Ghassemi et al. (Ghassemi, 2018) for an in-depth discussion of these features.

### 4.1 Complexity Features

Complexity features represent the degree of randomness and irregularity associated with the EEG signal. Different features in the form of entropy and complexity measures were extracted to gauge the information content of non-linear and non-stationary EEG signal data.

#### 4.1.1 Shannon Entropy

Shannon entropy (Shannon, 1948) is a measure of uncertainty (or variability) associated with a random variable. Let X be a set of finite discrete random variables *X* = {*x*_1_, *x*_2_, …, *x*_*m*_}, *x*_*i*_ ∈ *R*^*d*^, Shannon entropy, *H*(*X*), is defined as

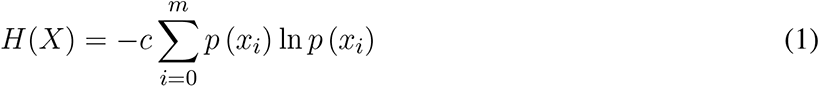

where c is a positive constant and p(*x*_*i*_) is the probability of (*x*_*i*_) (*ϵ*) X such that:

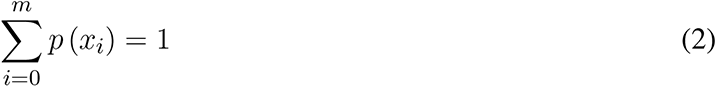

Higher values of entropy are indicative of high complexity and less predictability in the system. (Phung et al., 2014)

#### 4.1.2 Subband Information Quantity

Sub-band Information Quantity (SIQ) refers to the entropy of the decomposed EEG wavelet signal for each of the five frequency bands.(Jia et al., 2008; Valsaraj et al., 2020). In our analysis, the EEG signal was decomposed using a butter-worth filter of order 7 followed by an FIR/IIR filter. Shannon entropy (*H*(*X*)) of this resultant wave signal is the desired SIQ of a particular frequency band. Due to its tracking capability for dynamic amplitude change and frequency component change, this feature has been used to measure the information contained in the brain (Shin et al., 2006; Kanungo et al., 2021).

#### 4.1.3 Hjorth Parameters

Hjorth Parameters indicate time-domain statistical properties introduced by Bo Hjorth in 1970 (Hjorth, 1970). Variance-based calculation of Hjorth parameters incurs a low computational cost which makes them appropriate for performing EEG signal analysis. We make use of complexity and mobility (Das and Pachori, 2021) parameters in our analysis. Horjth mobility signifies the mean frequency or the proportion of standard deviation of the power spectrum. It is defined as :

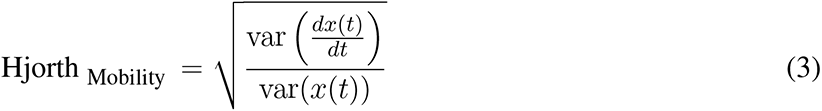

where var(.) denotes the variance operator and *x*(*t*) denotes the EEG time-series signal.

Hjorth complexity signifies the change in frequency. This parameter has been used to get a measure of similarity of the signal to a sine wave. It is defined as:

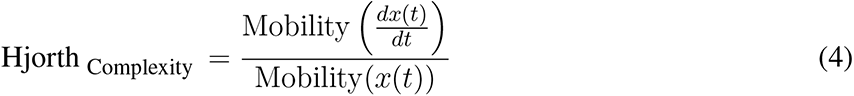

#### 4.1.4 False Nearest Neighbour

False Nearest Neighbour is a measure of signal continuity and smoothness. It is used to quantify the deterministic content in the EEG time series data without assuming chaos(Kennel et al., 1992; Hegger and Kantz, 1999).

#### 4.1.5 Asymmetry features

We incorporate Differential Entropy (DE) (Zheng et al., 2014) in our analysis to construct two features for each of the five frequency bands, namely, Differential Asymmetry (DASM)and Rational Asymmetry (RASM). Mathematically, DE (*h*(*X*)) is defined as :

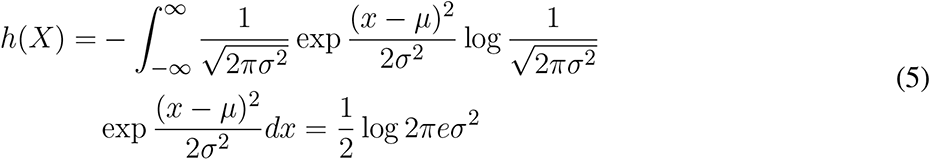

where *X* follows the Gauss distribution *N* (*µ, σ*^2^), *x* is a variable and *π* and exp are constant.

Differential Asymmetry(or DASM) (Duan et al., 2013) for each frequency band were calculated as the difference of differential entropy of each of 7 pairs of hemispheric asymmetry electrodes.

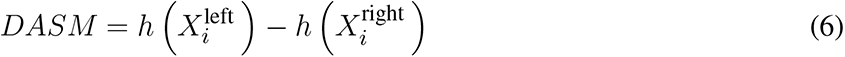

Rational Asymmetry(or RASM) (Duan et al., 2013) for each frequency band were calculated as the ratio of differential entropy between each of 7 pairs of hemispheric asymmetry electrodes.

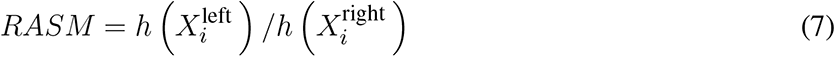

### 4.2 Continuity Features

Continuity features signify the clinically relevant signal characteristics of EEG signals(Hirsch et al., 2013; Ghassemi, 2018). These features have been acclaimed to serve as qualitative descriptors of states of the human brain and hence, are important towards the process of emotion recognition.

#### 4.2.1 Median Frequency

Median Frequency refers to the 50% quantile or median of the power spectrum distribution. Median Frequency has been studied extensively in the past due to its observed correlation with awareness (Schwilden, 1989) and its ability to predict imminent arousal(Drummond et al., 1991). It is a frequency domain or spectral domain feature.

#### 4.2.2 Band Power

Band power refers to the average power of the signal in a specific frequency band. The band powers of delta, theta, alpha, beta, and gamma were used as spectral features. To calculate band power, initially, a butter-worth filter of order 7 was applied on the EEG signal. IIR/FIR filter was applied further on the EEG signal in order to separate out signal data corresponding to a specific frequency band. Average of the power spectral density was calculated using a periodogram of the resulting signal. Signal Processing sub module (scipy.signal) of SciPy library (Virtanen et al., 2020) in python was used to compute the band power feature.

#### 4.2.3 Standard Deviation

Standard Deviation has proved to be an important time-domain feature in the past experiments (Amin et al., 2017; Panat et al., 2014). Mathematically, it is defined as the square root of variance of EEG signal segment.

#### 4.2.4 Diffuse Slowing

Previous studies (Boutros, 1996) have shown that diffuse slowing is correlated with impairment in awareness, concentration, and memory and hence, it is an importance feature for estimation of valence/arousal levels from EEG signal data.

#### 4.2.5 Spikes

Spikes(Hirsch et al., 2013) refer to the peaks in the EEG signal up to a threshold, fixed at mean + 3 standard deviation. The number of spikes was computed by finding local minima or peaks in EEG signal over 7 samples using scipy.signal.find peaks method from SciPy library (Virtanen et al., 2020).

#### 4.2.6 Delta Burst after spike

The change in delta activity after and before a spike computed epoch wise by adding mean of 7 elements of delta band before and after the spike, used as a continuity feature.

#### 4.2.7 Sharp spike

Sharp spikes refer to spikes which last less than 70ms and is a clinically important feature in study of electroencephalography (Hirsch et al., 2013).

#### 4.2.8 Number of Bursts

The number of amplitude bursts(or simply number of bursts) constitutes a significant feature (Hirsch et al., 2013).

#### 4.2.9 Burst length mean and standard deviation

Statistical properties of the bursts, mean *µ* and standard deviation *σ* of the burst lengths, have been used as continuity features.

#### 4.2.10 Number of Suppressions

Burst Suppression refers to a pattern where high voltage activity is followed by an inactive period and is generally a characteristic feature of deep anaesthesia(Ching et al., 2012). We use the number of contiguous segments with amplitude suppressions as a continuity feature with a threshold fixed at 10*µ* (Saba-Sadiya *et al*., *2020)*.

#### 4.2.11 Suppression length mean and standard deviation

*Statistical properties like mean µ* and standard deviation *σ* of the suppression lengths, used as a continuity feature.

## 5 FEATURE SELECTION

Selecting the correct predictor variables or feature vectors can improve the learning process in any machine learning pipeline. In this work, initially, sklearn’s (Pedregosa et al., 2011) VarianceThreshold feature selection method was used to remove zero-variance or constant features from the set of 36 extracted EEG features. Next, a subset of 25 features common to all 3 datasets (DREAMER, DEAP, and OASIS) was selected after applying the VarianceThreshold method for further analysis. This was done to validate our approach on a common set of features. The set of 11 features (S.E., F.N.N., D.S., S.K., D.B.A.S., N.B., B.L.M., B.L.S., N.S., S.L.M., S.L.S.) were excluded from further analysis. In our study, SelectKBest is used as a feature ranking and selection technique for all 3 datasets.

SelectkBest (Pedregosa et al., 2011) is a filter-based, univariate feature selection method intended to select and retain first k-best features based on the scores produced by univariate statistical tests. In our work, f regression was used as the scoring function since valence and arousal are continuous numeric target variables. It uses Pearson correlation coefficient as defined in Eq 8 to compute the correlation between each feature vector in the input matrix, X and target variable, y as follows:

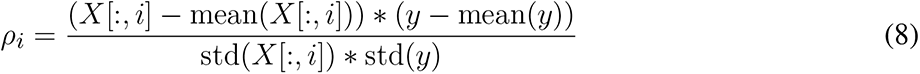

The corresponding F-value is then calculated as:

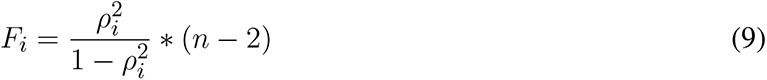

where n is the number of the samples.

SelectkBest method then ranks the feature vectors based on F-scores returned by f regression method. Higher scores correspond to better features.

## 6 RESULTS

### 6.1 Electrodes ranking and selection

The electrodes were ranked for the three datasets, using the SelectKBest method, as discussed in Section 5 and the ranks are tabulated for valence and arousal labels in Table 2. To produce a ranking for Top *N* electrodes taken together, feature data for top i electrodes were initially considered. The resultant matrix was split in the ratio 80:20 for training and evaluation of the random forest regressor model. The procedure was repeated until all the 14 electrodes were taken into account. The RMSE values for the same are shown in Fig 4 (a). It should be noted that, unlike feature analysis, data corresponding to 5 features each of DASM and RASM was excluded from the Top N electrode-wise RMSE study since these features are constructed using pairs of opposite electrodes.

**Table 1.** Table summarising various machine learning algorithms and features used to classify emotions on various datasets, reported with accuracy

| Dataset | Classifier | Feature extraction method | Accuracy (%) | Ref |
| --- | --- | --- | --- | --- |
| DEAP | kNN | Gray-Level Co-occurrence Matrix; Spectral Power Density | 79.58 (Average) | (Jadhav et al., 2017) |
| DEAP | kNN | Relative Power Energy; Logarithmic Relative Power Energy; Absolute Logarithmic; Relative Power Energy | 67.51, 68.55, 65.10 (VAD) | (Verma and Tiwary, 2017) |
| DEAP | SVM | Hjorth parameters; Entropy; Power of Frequency bands; RASM; DASM; Energy of frequency bands using wavelets | 65.72 (10 fold CV); 65.92 (LOO-CV) | (Khateeb et al., 2021) |
| DEAP | GNB | Spectral power; Spectral power differential asymmetry | 61.6, 64.7, 61.8 (VAD) | (Koelstra et al., 2012) |
| Video Clips | kNN | Absolute logarithmic Recoursing; Energy Efficiency of alpha, beta, and gamma bands decomposed using db4 wavelet function | 83.26 | (M et al., 2010) |
| Movie Clips | SVM | Power spectrum and wavelet decomposition of frequency bands; Entropy exponent; Katz fractal dimension; Feature smoothening using LDS; Feature reduction using PCA, LDA and CFS | 87.53 (Best accuracy) | (Wang et al., 2011a) |
| DEAP | 2k-NN | Spectral power of frequency bands; Spectral power difference of symmetric electrodes; Histogram parameters of segment level probability vectors; Dirichlet distribution parameters | 76.9, 68.4, 73.9, 75.3 (VADL) | (Wang et al., 2014) |

**Table 2.** Electrode Ranking for valence label (V) and arousal label (A) based on SelectKBest feature selection method.

| Electrode | DREAMER |  | DEAP |  | OASIS |  |
| --- | --- | --- | --- | --- | --- | --- |
|  | A | V | A | V | A | V |
| AF3 | 10 | 13 | 10 | 8 | 7 | 6 |
| AF4 | 9 | 11 | 12 | 10 | 8 | 8 |
| F3 | 11 | 10 | 7 | 11 | 5 | 5 |
| F4 | 13 | 14 | 8 | 6 | 6 | 9 |
| F7 | 14 | 12 | 1 | 1 | 1 | 1 |
| F8 | 3 | 5 | 2 | 2 | 4 | 4 |
| FC5 | 5 | 9 | 14 | 14 | 10 | 7 |
| FC6 | 6 | 4 | 3 | 4 | 9 | 10 |
| O1 | 12 | 8 | 13 | 7 | 11 | 11 |
| O2 | 8 | 3 | 6 | 9 | 14 | 13 |
| P7 | 4 | 2 | 5 | 3 | 12 | 12 |
| P8 | 7 | 6 | 4 | 5 | 13 | 14 |
| T7 | 1 | 7 | 9 | 13 | 3 | 2 |
| T8 | 2 | 1 | 11 | 12 | 2 | 3 |

**Figure 4.**
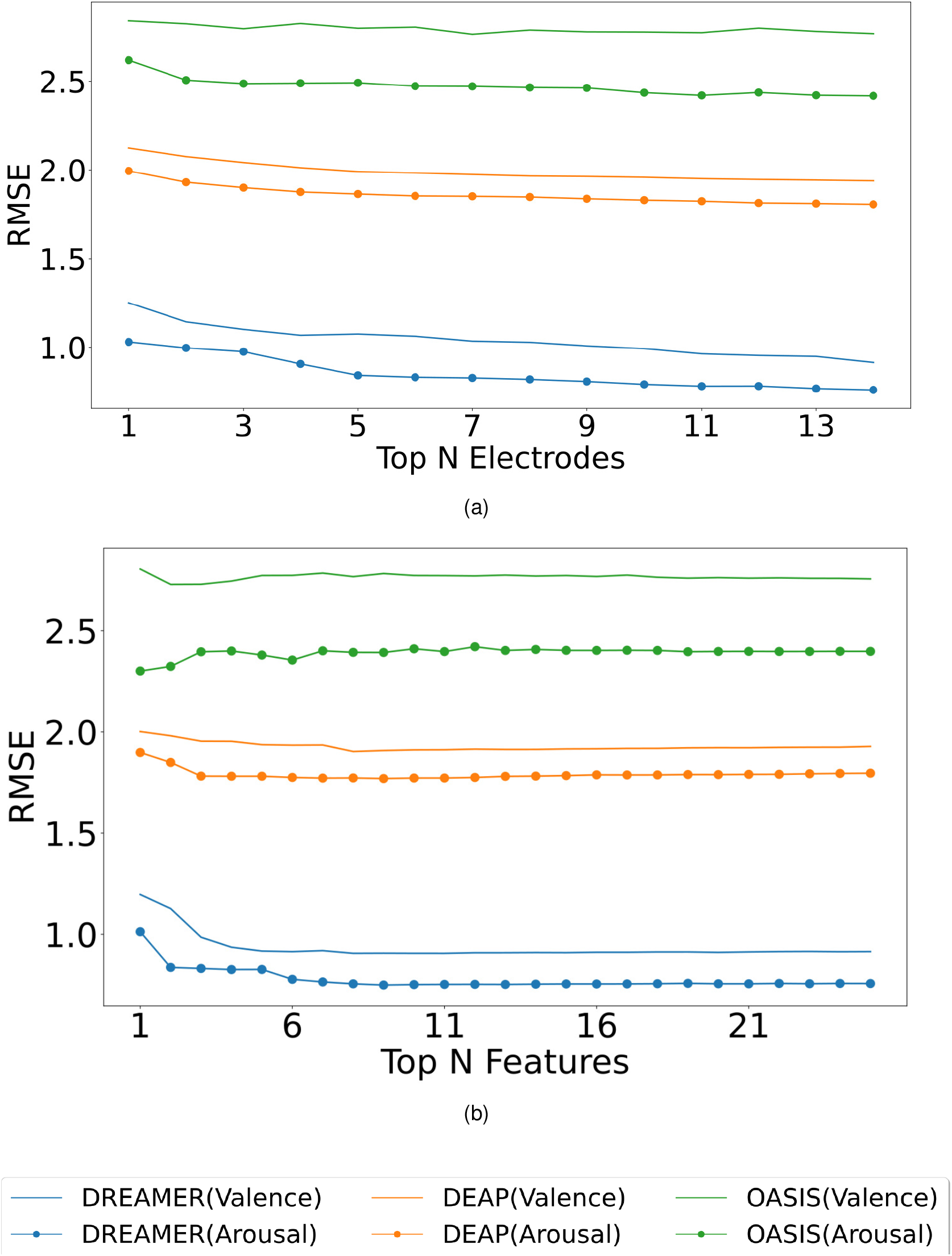
Model evaluation for feature and electrode selection. The random forest regressor was trained on the training set (80%) corresponding to top N electrodes (ranked using SelectKBest feature selection method) and RMSE was computed on the test set (20%) for valence (plain) and arousal (dotted) label on DREAMER, DEAP and OASIS EEG datasets as shown in (a). A similar analysis was performed for top N features for DREAMER, DEAP and OASIS EEG datasets as shown in (b).

### 6.2 Features ranking and selection

Each extracted feature was used to generate its corresponding feature matrix of shape (nbChannels, nbSegments). These feature matrices were then ranked using SelectKBest feature selection method. Initially, a feature matrix for the best feature was generated. The ranks were tabulated for valence and arousal labels in Table 3. This data was split into 80:20 train-test data, the training data was used to perform regression with Random Forest Regressor and predicted values on test data were compared with actual test labels, and RMSE was computed. In the second run, feature matrices of best and second-best features were combined, data was split into train and test data, model was trained, and predictions made by model on test data were used to compute RMSE. This procedure was followed until all the features are taken into account. The RMSE values for the feature analysis procedure, as described above, are shown in Fig 4 (b).

**Table 3.** Feature Ranking for valence label(V) and arousal label (A) based on SelectKBest feature selection method

| Feature | DREAMER |  | DEAP |  | OASIS |  |
| --- | --- | --- | --- | --- | --- | --- |
|  | A | V | A | V | A | V |
| B.P.A. | 7 | 5 | 18 | 17 | 25 | 15 |
| B.P.B. | 9 | 8 | 8 | 7 | 11 | 13 |
| B.P.D. | 22 | 22 | 23 | 22 | 12 | 23 |
| B.P.G. | 21 | 18 | 3 | 13 | 5 | 20 |
| B.P.T. | 20 | 20 | 19 | 18 | 24 | 17 |
| D.A. | 4 | 11 | 12 | 10 | 16 | 9 |
| D.B. | 12 | 10 | 4 | 4 | 21 | 11 |
| D.D. | 24 | 25 | 16 | 16 | 20 | 21 |
| D.G. | 16 | 16 | 5 | 6 | 14 | 18 |
| D.T. | 13 | 14 | 10 | 9 | 23 | 16 |
| H.C. | 2 | 4 | 20 | 20 | 4 | 4 |
| H.M. | 6 | 3 | 17 | 19 | 1 | 2 |
| M.F. | 14 | 12 | 24 | 25 | 7 | 5 |
| R.A. | 5 | 13 | 21 | 21 | 15 | 12 |
| R.B. | 11 | 9 | 1 | 2 | 19 | 10 |
| R.D. | 23 | 24 | 25 | 24 | 18 | 22 |
| R.G. | 17 | 17 | 2 | 1 | 13 | 19 |
| R.T. | 15 | 15 | 11 | 11 | 22 | 14 |
| S.E.A. | 10 | 7 | 13 | 12 | 10 | 24 |
| S.E.B. | 3 | 2 | 6 | 5 | 3 | 3 |
| S.E.D. | 18 | 19 | 15 | 15 | 8 | 7 |
| S.E.G. | 1 | 1 | 9 | 8 | 2 | 1 |
| S.E.T. | 19 | 21 | 14 | 14 | 9 | 8 |
| S.S.N. | 25 | 23 | 22 | 23 | 17 | 25 |
| S.D. | 8 | 6 | 7 | 3 | 6 | 6 |

The identification of an optimum set of electrodes and features is a critical step. By optimum set, we imply the minimum number of electrodes and features that produce minimum RMSE during model evaluation, as shown in Fig 4. We can observe a general decline in RMSE value when the number of electrodes under consideration is increased. DREAMER dataset shows a much greater and smoother convergence than the other two datasets because more training data was available for training the model. In general, the minimum RMSE is observed when all 14 electrodes are selected. OASIS dataset can be excluded from this inference since it contained only 15 participants at the time of the experiment.

Fig 4 (b), reveals a general pattern about optimal set of features. On increasing the number of features in consideration, initially, there is a steady drop in the RMSE values followed by a gradual increase after a certain critical point in the graph. Hence, a minima can be observed in the graph. As discussed above, the OASIS dataset can be ruled out of this generalization. The minimum RMSE values and the corresponding number of best features and electrodes selected are summarized in Table 4 and Table 5 respectively.

**Table 4.**
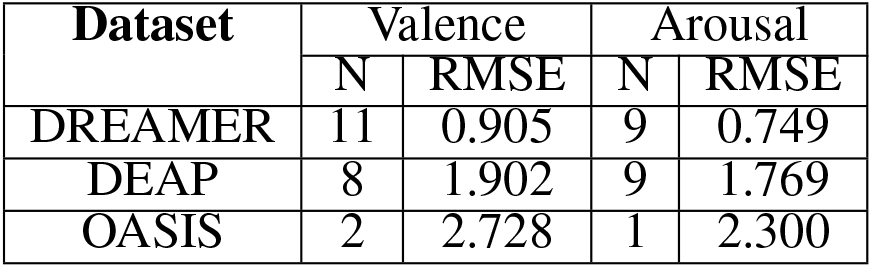
RMSE values for valence and arousal label on the test set (20%) of DEAP, DREAMER and OASIS dataset for optimum set of features.

**Table 5.**
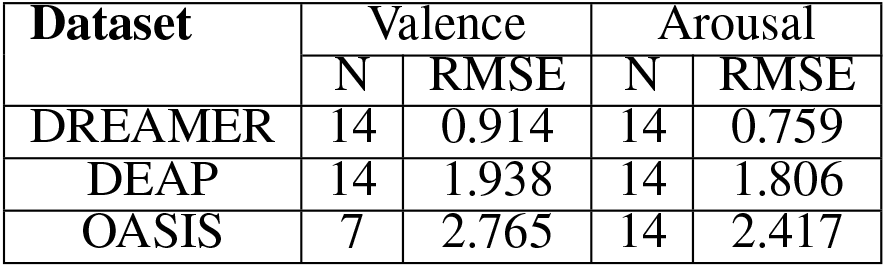
RMSE values for valence and arousal label on the test set (20%) of DEAP, DREAMER and OASIS dataset for optimum set of electrodes.

#### Incremental Learning

As given by the feature analysis described above, the best features were used to generate a feature matrix for valence and arousal for each dataset. The feature matrix was then used to train a random forest regressor as part of the incremental learning algorithm.

Incremental learning was performed based on the collection of subject data. Initially, the first subject data was taken, their trial order shuffled and then split using 80:20 train test size, the model was trained using train split, predictions were made for test data, next 2^*nd*^ subject data was taken together with the 1^*st*^ subject, trial order shuffled, again a train-test split taken and the random forest regressor model was trained using the train split. Predictions were made for the test split. This procedure was repeated until data of all the subjects were used for RMSE computation. RMSE values for each training step, i.e. training data consisted of subject 1 data, then the combination of subject 1, 2 data, then the combination of subject 1, 2, 3 data, and so on. The plots generated for RMSE values for the individual steps of training show a general decreasing trend as evident from Fig 5.

**Figure 5.**
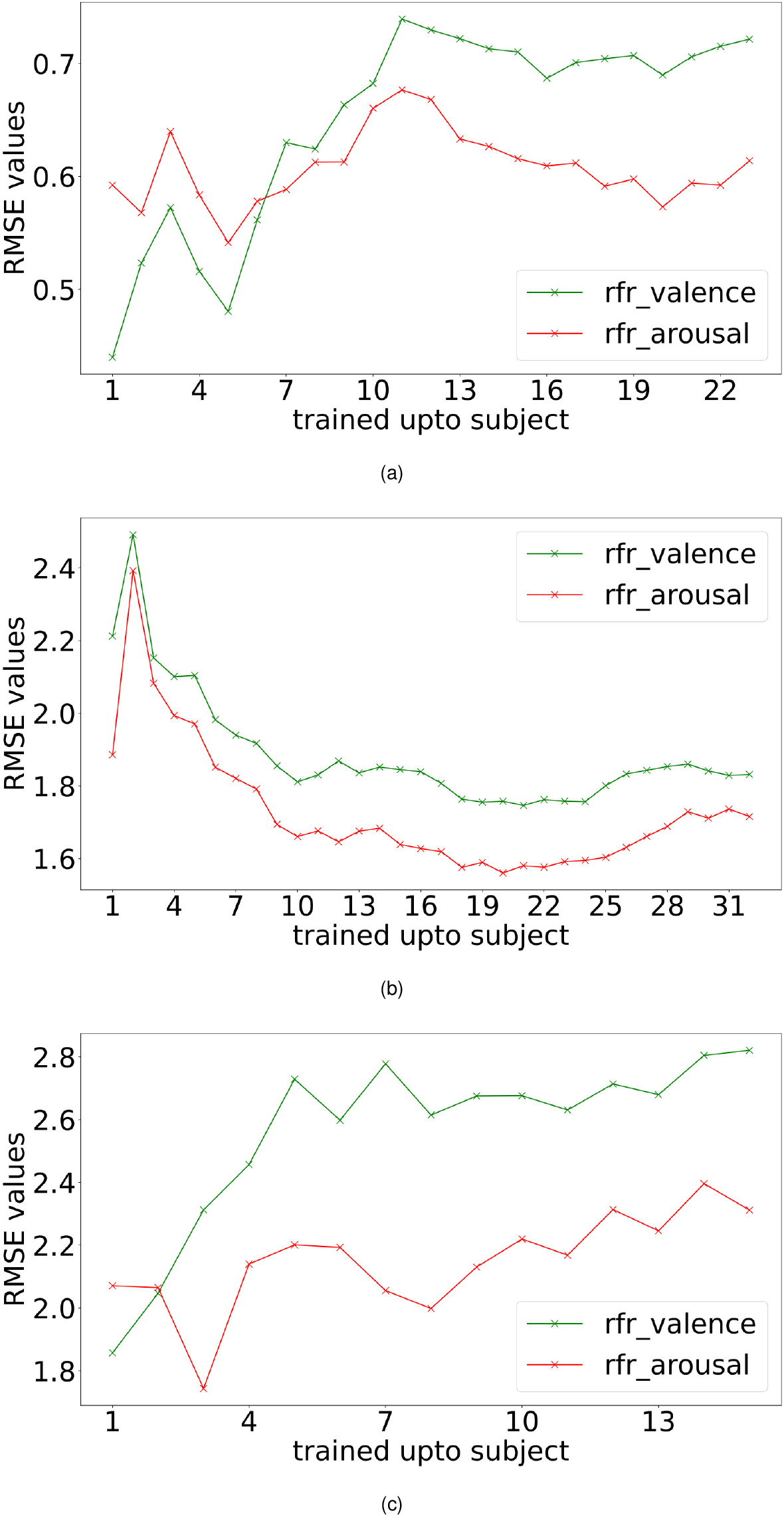
Incremental learning performance. Valence and arousal RMSE readings obtained with incremental learning for DREAMER (a), DEAP (b) and OASIS (c) EEG dataset using random forest regressor (rfr).

#### Leave-one-subject-out cross-validation

Subject generalization is a crucial problem in identifying EEG signal patterns. To prevent over-fitting and avoid subject-dependent patterns. We train the model with data of all the subjects except a single subject and evaluate the model on this remaining subject. Hence, the model is evaluated for each subject to identify subject bias and prevent any over-fitting. Also, when building a machine learning model, it is a standard practice to validate the results by leaving aside a portion of data as the test set. In this work, we used the leave-one-subject-out cross-validation technique due to its robustness for validating results for data set at the participant level. Leave-one-subject-out cross-validation is a k-fold cross-validation technique, where the number of folds, k, equals the number of participants in a dataset. The cross-validated RMSE values for the three datasets for all the participants are plotted in Fig 6.

**Figure 6.**
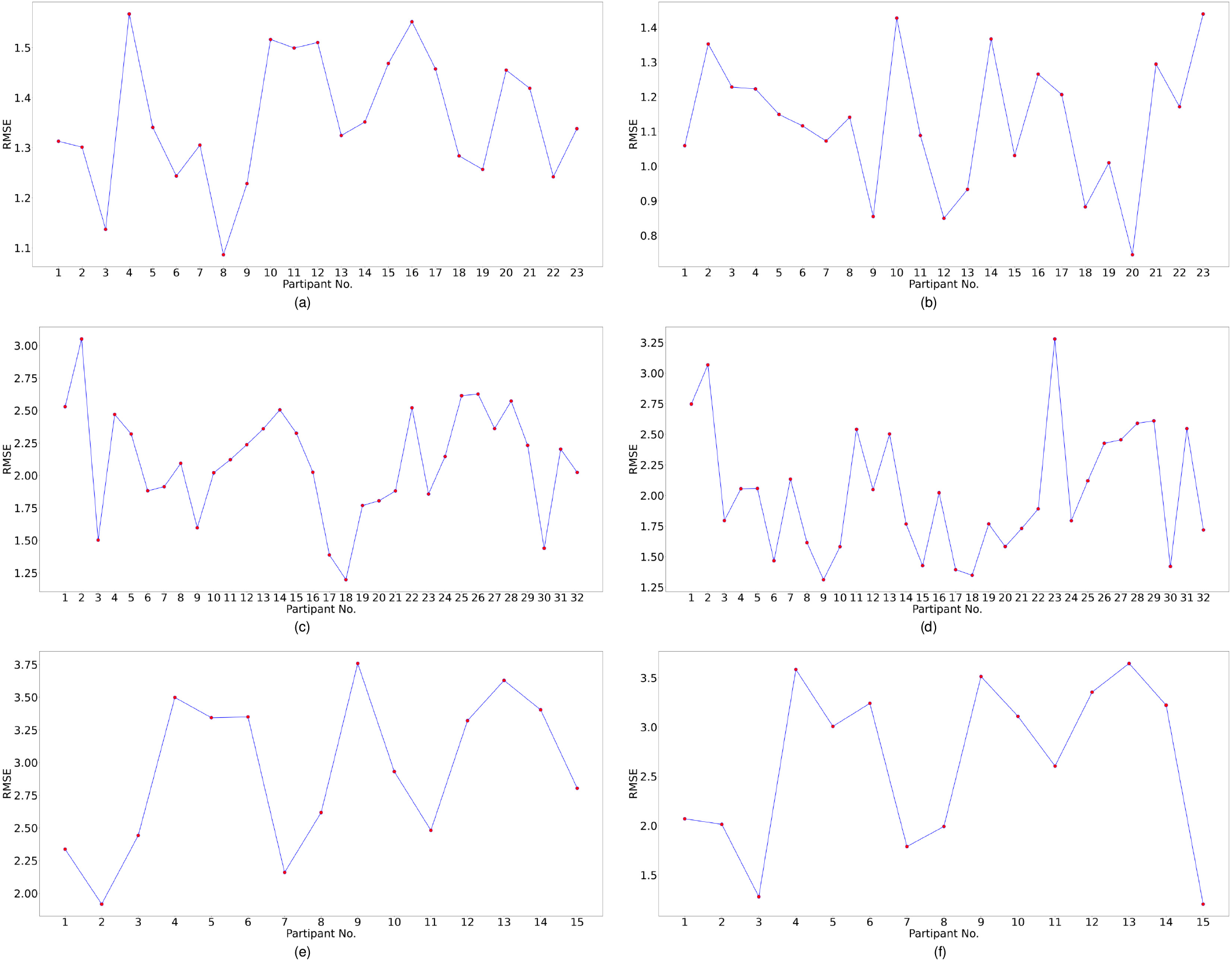
Subject wise performance analysis for valence and arousal labels. Leave-one-subject-out cross-validation performance analysis for valence label for (a) DREAMER (b) DEAP (c) OASIS datasets and for arousal label for (d) DREAMER (e) DEAP (f) OASIS datasets respectively. In this cross-validation technique, one subject was chosen as the test subject, and the models were trained on the data of the remaining subjects.

Mean and standard deviation of RMSE values for valence and arousal label after cross validation have been summarized in Table 6. The best RMSE values lie within the standard deviation range respect to the leave-one-subject-out cross validation results and hence, inferences drawn from them can be validated.

**Table 6.**
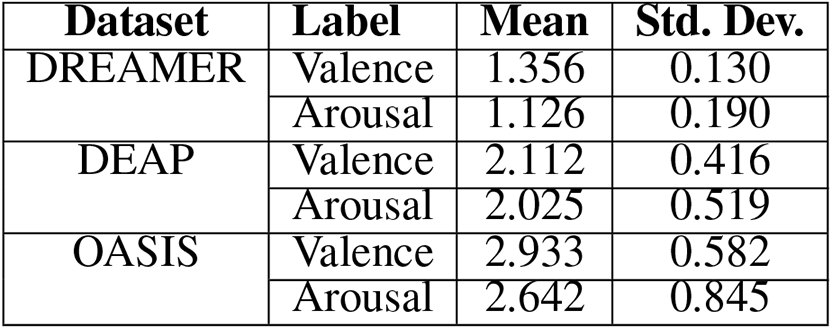
Mean and Standard Deviation (Std. Dev.) of RMSE values for Valence and Arousal Label Data after Leave-one-subject-out-cross-validation

## 7 DISCUSSION

The relation between the performance (RMSE) and the number of participants is critical for any study concerning emotion recognition from the EEG dataset. As in Fig 5, we observe an improvement in performance with an increasing number of participants. This explains that the machine learning algorithm needs data from more participants for generalization. Interestingly, the performance degrades for the OASIS dataset (Fig 5 (c)) while increasing the number of participants. This could be explained as the model overfits when trained with data from a few subjects. This can be verified by the fact that the degradation in performance is only up to a certain number of subjects, as in Fig 5 (a). Hence, with data from more participants in the OASIS EEG dataset, we can expect to observe an increase in performance.

As shown in tables 2 and 3, 3 rankings were obtained as a result of 3 datasets for each label. For the valence labels, out of the top 25 % electrodes, 33 % were in the frontal regions (F3, F4, F7, F8, AF3, AF4, FC5, FC6), 33% in the temporal regions (T8, T7), 22 % the parietal regions (P7, P8), and 11 % in the occipital regions (O1, O2). For the top 50% electrodes, 57 % were in the frontal regions, 19 % in the temporal regions, 19 % in the parietal regions, and 4 % in the occipital regions.

For the arousal labels, out of the top 25 % electrodes, 55 % were in the frontal regions and 44 % in the temporal regions. For top 50% electrodes, 57 % were in the the frontal regions, 19 % in the temporal regions, 19 % in the parietal regions, and 4 % in the occipital regions.

Therefore, the frontal region was the most significant brain region for recognizing valence and arousal, followed by temporal, parietal, and occipital. This is in accordance with previous works on EEG channel selection (Alotaiby et al., 2015), (Shen et al., 2020a).

The optimum set of features for the DREAMER dataset was observed to be (S.E.G, S.E.B, H.M, H.C., B.P.A, S.D, S.E.A, B.P.B, R.B, D.B, D.A) for valence and (S.E.G, H.C, S.E.B, D.A, R.A, H.M, B.P.A, S.D, B.P.B) for arousal respectively. The minimum RMSE values obtained using these optimal features on the DREAMER dataset were 0.905 and 0.749 for valence and arousal dimensions, respectively, as evident from Table 4. Therefore these features were critical for recognizing emotional states and can be used in future studies to evaluate classifiers like Artificial Neural Networks and ensembles.

As shown in Table 3, band power and sub-band information quantity features for gamma and beta frequency bands performed better in the estimation of both valence and arousal than other frequency bands. Hence the gamma and beta frequency bands are the most critical for emotion recognition (Wang et al., 2011b), (Zheng et al., 2017).

It can be inferred from Tables 3 that H.M. was mostly ranked among the top 3 features for predicting valence labels and arousal labels. Similarly, H.C. was ranked among the top 4 features. This inference is consistent with the previous studies that claim the importance of time-domain Hjorth parameters in accurate EEG classification tasks (Türk et al., 2017; Cecchin et al., 2010).

In the past, statistical properties like standard deviation derived from the reconstruction of EEG signals have been claimed to be significant descriptors of the signal and provide supporting evidence to the results obtained in this study (Malini and Vimala, 2016; Panda et al., 2010). It was observed that SD was ranked among the top 8 ranks in general.

Table 4 indicates that the minimum RMSE values obtained on the test set (20%) using the optimum set of features were 0.905 and 0.749 on the DREAMER dataset, 1.902 and 1.769 on the DEAP dataset and 2.728 and 2.3 on OASIS dataset for valence and arousal respectively. For leave-one-subject-out cross-validation, we achieved the best RMSE of 1.35, 1.126 on DREAMER, 2.11, 2.02 on DEAP and 2.93, 2.64 on the OASIS dataset for valence and arousal, respectively as shown in Fig 6.

## 8 CONCLUSION AND FUTURE SCOPE

EEG is a low-cost, noninvasive neuroimaging technique that provides high spatiotemporal information about brain activity, and it has become an indispensable tool for decoding cognitive neural signatures. However, the multi-stage intelligent signal processing method has several indispensable steps like pre-processing, feature extraction, feature selection, and classifier training. In this work, we propose a generalized open-source neural signal processing pipeline based on machine learning to accurately classify emotional index on a continuous valence-arousal plane using these EEG signals. We statistically investigated and validated artifact rejection, automated bad-trial rejection, state-of-the-art Spatio-temporal feature extraction techniques, and feature selection techniques on a self-curated dataset recorded from a portable headset in response to OASIS emotion elicitation image dataset and two open source EEG datasets. This published dataset could be used in future studies for a spectrum of intelligent signal processing methods like deep learning, reinforcement learning, and neuromorphic computing. The published simplistic python pipeline would aid researchers in focusing on innovation in specific signal processing steps like feature selection or machine learning without the need to recreate the entire pipeline from scratch. In accordance with neuroscience literature, our proposed system could identify the optimum set of electrodes and features that produce minimum RMSE during emotion classification for a given dataset. It also validated the claim that beta and gamma frequency bands are more effective than other bands when it comes to emotion classification. We performed the evaluation of EEG activity induced by videos (DEAP), and static images (DREAMER & OASIS), but not on audio stimulus. The OASIS dataset collection was limited to 15 participants due to the Covid-19 pandemic. In future we plan to collect the data for at least 40 participants to draw stronger inferences. Future work would also include analysis of end to end neural networks and transfer learning for the purpose of emotion recognition. The published dataset can be used for further advancement of machine learning systems for emotional state detection with a data recorded from portable headset. The published EEG processing pipeline of artifact rejection, feature extraction, feature ranking, feature selection and machine learning could be expanded and adapted for processing EEG signal in response to variety of stimuli.

## AUTHOR CONTRIBUTIONS

N.G. conceptualized the research. R.G., N.G., A.A. performed the experiments and analyzed the data. V.B. supervised the study. All authors approved and contributed in writing the manuscript.

## ACKNOWLEDGMENTS

We acknowledge Mr. Parrivesh N.S. and Mr. V.A.S. Abhinav for their valuable suggestions and assistance in data collections during the planning and initial development of this research work. We acknowledge the support from Ironwork Insights Inc.

This work was supported by the Department of Science and technology, Government of India, vide Reference No:SR/CSI/50/2014(G)through the Cognitive Science Research Initiative (CSRI). We acknowledge financial supports from the EU: ERC-2017-COG project IONOS (GA 773228)

## DATA AVAILABILITY STATEMENTS

The dataset recorded in this study would be made available in public domain upon acceptance of manuscript. The code repository developed would be published at: https://github.com/rohitgarg025/Decoding_EEG

